# Uncovering and engineering the mechanical properties of the adhesion GPCR ADGRG1 GAIN domain

**DOI:** 10.1101/2023.04.05.535724

**Authors:** L. Dumas, M. Marfoglia, B. Yang, M. Hijazi, A.N. Larabi, K. Lau, F. Pojer, M.A. Nash, P. Barth

## Abstract

Key cellular functions depend on the transduction of extracellular mechanical signals by specialized membrane receptors including adhesion G-protein coupled receptors (aGPCRs). While recently solved structures support aGPCR activation through shedding of the extracellular GAIN domain, the molecular mechanisms underpinning receptor mechanosensing remain poorly understood. When probed using single-molecule atomic force spectroscopy and molecular simulations, ADGRG1 GAIN dissociated from its tethered agonist at forces significantly higher than other reported signaling mechanoreceptors. Strong mechanical resistance was achieved through specific structural deformations and force propagation pathways under mechanical load. ADGRG1 GAIN variants computationally designed to lock the alpha and beta subdomains and rewire mechanically-induced structural deformations were found to modulate the GPS-Stachel rupture forces. Our study provides unprecedented insights into the molecular underpinnings of GAIN mechanical stability and paves the way for engineering mechanosensors, better understanding aGPCR function, and informing drug-discovery efforts targeting this important receptor class.

---

Cellular mechanosensing is currently thought to primarily involve integrins, which couple the extracellular matrix (ECM) to the contractile cytoskeleton (*1*–*3*), or mechano-activated ion channels (*4*, *5*). However, adhesion GPCRs (aGPCRs) bearing large extracellular regions (ECR) can bind to ECM components (*6*–*8*) and have received growing attention in mechanical regulation (*9*–*13*). Virtually all receptors in this family bear a highly conserved GPCR-Autoproteolysis INducing (GAIN) domain located in the ECR, between the adhesion ligand binding domain (LBD) and the signaling 7 transmembrane (7TM) domain. The GAIN structure carries a tethered peptide agonist (TA or Stachel) that often undergoes auto-proteolytic cleavage but remains deeply buried inside the GAIN structure (*6*, *14*). While several lines of evidence support an allosteric mechanism of receptor activation that does not involve Stachel dissociation (*6*, *15*–*18*), aGPCR signaling under mechanical stress can also occur upon Stachel dissociation from GAIN. With GAIN dislodged, Stachel then binds the 7TM domain and triggers intracellular signaling (*7*, *15*, *17*). Recent cryoEM structures of aGPCRs in active signaling states (i.e. bound to GAIN-dissociated TA and G-proteins) support this receptor activation mechanism through GAIN shedding (*19*–*22*). However, reconciling the allosteric and shedding mechanisms of activation has been challenging, and would necessitate an understanding of the molecular underpinnings of GAIN-mediated mechanotransduction, which have remained elusive so far.

Protein structures undergo specific deformation upon mechanical loading which guides force-activated phenotypes and resistance to applied stress (*23*, *24*). In particular, protein topology and the orientation of chemical bonds with respect to the direction of the mechanical force propagating inside the structure are key determinants of protein mechanical responses (*23*–*25*). In recent years, proteins with novel shapes and topologies have been designed *de novo* by optimizing folding and thermodynamic stability in the absence of perturbations (i.e. under equilibrium conditions) (*26*). Engineering protein stability under mechanical stress would enable the design of powerful mechano-sensing functions but this approach remains poorly exploited in protein design.

In this study, we aimed to uncover the mechanical properties of the GAIN domain. We selected and studied a prototypical GAIN domain, that of the widely studied ADGRG1 receptor, using single-molecule atomic-force spectroscopy (AFM) and steered molecular dynamics (SMD) simulations. To probe the mechanical unfolding observed by SMD, we computationally designed GAIN variants locked in specific conformational states. Designed GAINs exhibited altered mechanical stability in good agreement with our predictions (**Fig.1A**).

**Fig. 1.**
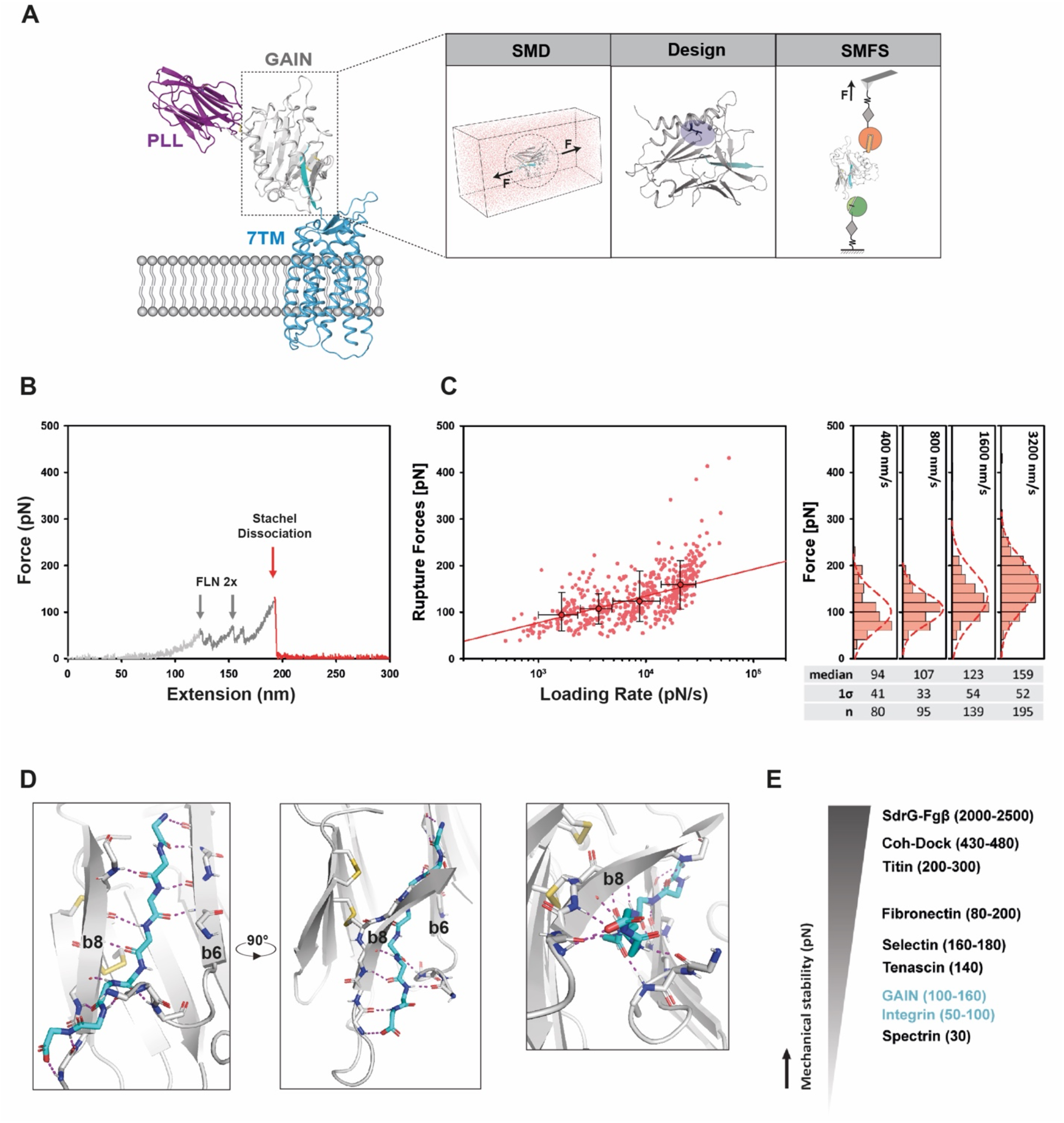
The adhesion GPCR GAIN-Stachel interface behaves as a stable mechanical clamp motif. (**A**) General framework for the study of the ADGRG1 GAIN domain under force. Representation of membrane-embedded full-length human ADGRG1 adapted from an AlphaFold structure prediction (UniProt Q9Y653), composed of the 7TM (blue), GAIN (grey, with Stachel in cyan) and PLL (magenta) domains (left panel). A combination of experimental and computational techniques was used to probe the behavior of WT and mutant GAIN domains under force (right insets). (**B**) Representative single-molecule force-extension trace of the WT ADGRG1 GAIN domain showing two FLN fingerprint unfolding events and a single GAIN-Stachel dissociation event at ∼200 nm extension. (**C**) Dynamic force spectrum of the GAIN domain at cantilever retraction velocities of 0.4 μm.s^-1^, 0.8 μm.s^-1^, 1.6 μm.s^-1^ and 3.2 μm.s^-1^, with corresponding rupture-force probability density histograms for each velocity projected onto individual axes on the right. A Gaussian fit (dashed line) with the fit parameters are shown for each distribution. (**D**) Structural view of the Stachel backbone (cyan) H-bond network (dotted magenta lines) formed with adjacent β-strands b6 and b8 in the human ADGRG1 GAIN domain. The right-most panel shows a view from C-terminal end of the Stachel. (**E**) Stability of human ADGRG1 GAIN-Stachel interaction situated within a spectrum of proteins (cyan: signaling proteins, black: adhesion proteins) involved in mechanical signaling and adhesion. Force values are indicative and given as broad reference points or ranges corresponding to the measured force at which a given protein or protein-protein interaction undergoes a major conformational transition important for its function (*23*, *33*, *36*, *40*– *42*). Values can vary depending on the experimental technique and setup used, the loading rate applied, the ligand used (if any) and the exact protein type and subdomain considered.

We selected the ADGRG1 receptor because it is a canonical and well-studied aGPCR that bears a topologically simple ECR composed of one N-terminal adhesion ligand-binding domain (PTX/LNS-Like, PLL) linked to the 7TM signaling region by an autoproteolytically cleaved GAIN domain. The structure of the isolated murine homolog ADGRG1 EC region was solved by X-ray crystallography (*27*). The cryoEM structure of the 7TM region in an active signaling conformation bound to the GAIN-dissociated TA and G-protein was released recently (*19*).

We first sought to measure the mechanical properties of the isolated GAIN domain using SMFS. It is known that a protein’s mechanical response is highly dependent on the loading point and direction of the applied force (*25*, *28*–*32*). In the context of the full-length ADGRG1 receptor expressed at the cell surface, mechanical force is likely transduced to GAIN by the N-terminal adhesion PLL domain bound to an ECM ligand. To best mimic this native orientation and direction of the applied force in our SMFS experiments, we designed a setup where the C-terminal end of GAIN was attached to a solid surface while the pulling force was applied to the N-terminal end of the domain (**Fig.1A**, **Methods**) through a regeneratable peptide/receptor complex (FgB/SdrG). This experimental design enabled fresh GAIN molecules to be probed on the surface and allowed repeated measurements and quantification of Stachel dissociation by AFM. We measured WT GAIN-Stachel dissociation events in the range of 100-160 pN with a clear dependence on the loading rate over a range from 1,000 – 40,000 pN/s (**Fig.1B-C**). These results indicate that the GAIN-Stachel interface is significantly more mechanically stable than other canonical signaling mechanosensors characterized using SMFS such as Notch and integrin receptors (**Fig.1E**) (*33*– *35*). To ensure that the observed events corresponded to the rupture of the GAIN-Stachel interface, we carried out AFM measurements on a non-autoproteolytically cleaved form of the GAIN bearing the T383G mutation. These traces uniformly exhibited high force rupture (∼2 nN) of the peptide handle/receptor (FgB/SdrG) with no structural unfolding transitions between the first FLN unfolding peak and the final rupture peak. Our data do not exclude the possibility of partial GAIN unfolding at low forces, however, they strongly suggest that dissociation is not driven by unfolding of GAIN (**Fig.S1**).

To elucidate which structural features might confer mechanostability to the GAIN-Stachel interface, we first analyzed the high-resolution X-ray structure of the murine ADGRG1 GAIN domain that shares high sequence homology to the human protein (*27*). The Stachel peptide adopts a β-strand conformation and interacts with two adjacent anti-parallel β-strands of GAIN through an extensive network of backbone hydrogen bonds (**Fig.1D**). These features are reminiscent of mechanical clamp motifs found in well-characterized Ig-like β-sandwich mechanical proteins of the human cardiac titins or bacterial adhesins for example (**Fig.1D**) (*31*, *39*–*41*). While in these systems, the hydrogen bond network holding the clamp contributes prominently to the protein domain’s overall force resistance, the mechanical stability of these interfaces can vary by up to two orders of magnitude. These observations suggest that the structural context, the overall topology of the fold, its deformation upon applied mechanical stress and how force propagates through the fold may also strongly determine mechanical resistance.

To better understand how the GAIN domain responds to mechanical force, we performed SMD simulations by applying a constant harmonic force potential to the C-terminal structure and pulling simultaneously from the N terminus of the GAIN to mimic the force directionality in AFM. We ran simulations at pulling speeds as low as 0.1 nm.ns^-1^. While this speed is orders of magnitude faster than those used in AFM, SMD performed at such rates were shown to recapitulate general unfolding and unbinding events measured by SMFS (*23*). Pulling on the protein from its N- and C-termini results in an increasing amount of force accumulated within the structure prior to unfolding or dissociation, and defines a loaded state of the system. We focused our analysis on the loaded state displaying the highest mechanical force resistance prior to Stachel dissociation (**Fig.2A**). In this state, GAIN undergoes topological rearrangements that favor its mechanical stability and reveal several key structural underpinnings of its mechanosensing properties. Specifically, we observed a rigid body reorientation of the N-terminal helices 1 and 2, a large conformational change of the loop connecting those helices and a partial unfolding of helix 1 (**Fig.2B,C**). Altogether, this structural reorganization created 3 polar interaction networks that were absent in the native structure (**Fig.2B,C**). These motifs stabilized the N-terminal part of the domain under mechanical stress by locking the 2 helices together with the first 2 β-strands of the GAIN scaffold. While the C-terminal β-strand region that buries the Stachel remained largely unperturbed in the loaded state, we observed subtle structural distortions and reorientation of the backbone hydrogen bonds stabilizing the Stachel within the GAIN. Overall these interactions adopted geometries with larger orthogonal components to the pulling force axis (**Fig.2D,E**), suggesting a stiffer and more mechanically resistant binding interface.

**Fig. 2.**
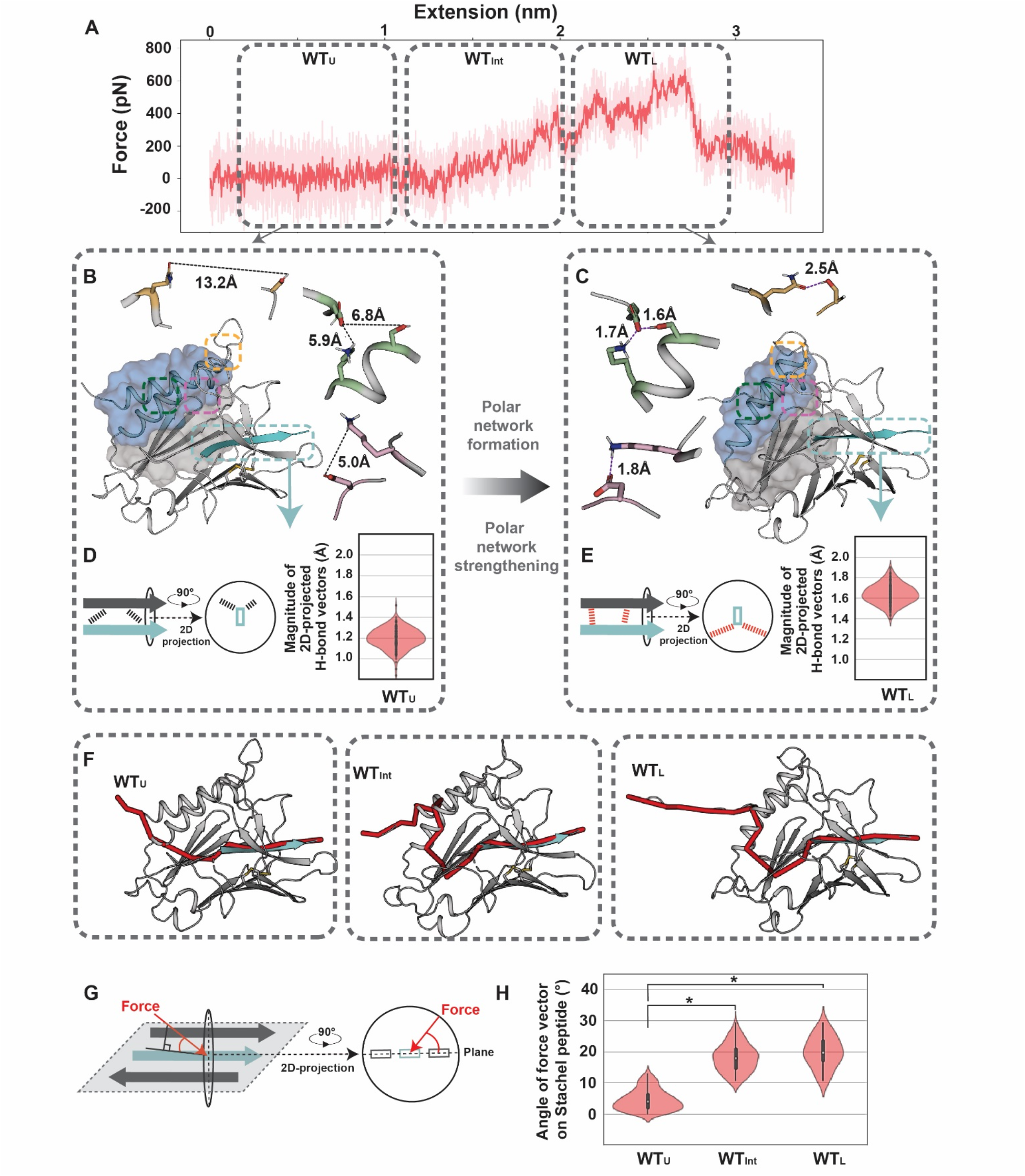
The GAIN domain structurally deforms and redirects the force propagation pathway upon mechanical loading. (A) Representative SMD trajectory (dark red) and rolling average (transparent red) leading to the loaded state. Gray dotted boxes delineate unloaded, intermediate and loaded regions of the SMD trajectories. **(B, C)** The GAIN N-terminal subdomain undergoes structural stabilization upon mechanical load through the formation of a strong non-native interaction network established between helices 1 and 2 (green box), loop 1 and helix 1 (yellow box), and helix 1 and β-strand 1-2 (pink box). These regions are highlighted on a GAIN structure where the N-terminal, C-terminal subdomains and Stachel are represented as a blue surface, grey surface and cyan cartoon, respectively. The residues forming each network are represented in sticks with corresponding contact distances. **(D, E)**. Magnitude of backbone hydrogen bond vectors projected on a plane orthogonal to the direction of the pulling axis (p-value: 0.0006). **(F)**. Force propagation pathways connecting the N- and C-termini of the GAIN in the unloaded (left), intermediate (middle) and fully loaded (right) states. **(G)**. Schematic view of the path angle calculation showing the force propagation vector and its associated angle. The Stachel (cyan) and interacting beta-strands (dark grey) are represented as cartoon. **(H)**. Angle distribution of force propagation pathways in the unloaded (WT_U_), partially loaded (WT_Int_) and fully loaded (WT_L_) states. *: p-value < 0.05.

To further understand how these structural deformations enable GAIN to withstand large mechanical loads, we analyzed how the applied force propagates through the structure and reaches the Stachel peptide. Several studies have shown that force propagation pathways through a system and the orientation of the force vector at the rupture interface strongly determine the system’s mechanical resistance (*25*, *43*–*45*). High mechanical resistance is achieved when the force propagates along pathways with strong components orthogonal to the pulling axis at the binding interface (*23*, *25*). We performed a correlation-based dynamical network analysis of the GAIN motions extracted from SMD to determine these specific paths. To extract the orientation of force on the Stachel mechanical clamp, the angles between the force pathway direction and the Stachel H-bond network plane were computed (**Fig.2G**).

In the absence of applied force, predicted paths correspond to the main long-range communication pipelines built in the native GAIN structure and are reminiscent of pathways found in allosteric proteins connecting regulatory and effector sites (*46*, *47*). As shown in **Fig.2F**, the unloaded pathway traverses the α-domain from the N-terminal helix 1 pulling point to the C-terminal of helix 2, then jumps onto β-strand 3 **(Fig.S3)** and travels through the β-domain to reach the Stachel with a direction almost colinear to the rupture interface (**Fig.2H**). By contrast, upon accumulation of mechanical stress in the loaded state, the main force propagation path changes and travels through helix 1 to reach the polar motif locking the 2 helices, jumps onto helix 2 and then β-strand 2 (**Fig.2F-H**) to reach the Stachel with an average angle of 19 degrees with respect to the rupture interface (**Fig.2J**). Our analysis indicates that the GAIN structure undergoes topological rearrangements that not only enhance the stability of local structures (e.g., the interface between the first 2 helices and β-strands, the H-bond network between the Stachel and neighboring β-strands) but also reorient the main force propagation path to reach the Stachel in a suboptimal direction for triggering peptide dissociation.

This analysis reveals major structural and dynamic underpinnings of GAIN’s mechanical properties. In particular, while distant from the Stachel, the α-domain is predicted to play a central role in the mechanical resistance of the rupture interface. Unlike the conserved β-domain, the sequence, structure and topology of the α-domain vary significantly between members of the aGPCR family (**Fig.S2**). This suggests that specific structural rearrangements and long-range communication of the force transmitted by the α-domain may significantly impact the precise mechanosensing properties of different GAIN domains.

To address this hypothesis, we attempted to fine tune the GAIN’s mechanical resistance by reengineering the α-β domain interface and rewiring these force propagation paths. Since large helical rigid body and loop motions occur upon mechanical load in our simulations, we reasoned that designing topological locks with disulfide bridges (*48*, *49*) that trigger or prevent such conformational changes should maximize the impact of sequence changes on force propagation and GAIN mechanical resistance, and stringently test our predictions. We first scanned the entire GAIN structure for Cα positions satisfying the formation of optimal disulfides and then selected a few designed variants with interesting predicted mechanical behavior (**Methods**).

We designed a first variant with a disulfide bridge (referred to SS_h2l10), which locks loop 10 connecting β-strands 8 and 9 onto the N-terminal of helix 2, and prevents the disruption of that α-β local interface observed when the WT GAIN reaches the loaded state. Unexpectedly, our calculations suggested that the force pathway for variant SS_h2l10 first propagates along helix 1 but then runs through the engineered disulfide onto the β-domain and reaches the Stachel, bypassing most of the residue network engaged in force propagation in the WT (**Fig.3A-C**). This rewired pathway reached the Stachel rupture interface at much more orthogonal angles than WT (69° and 63° compared to 19° and 36° for the WT optimal path at 1.0 and 0.1 nm.ns^-1^; **Fig.3J**). Hence, our model predicted higher mechanical stability of the GAIN-Stachel interface in the SS_h2l10 compared to the WT.

**Fig. 3.**
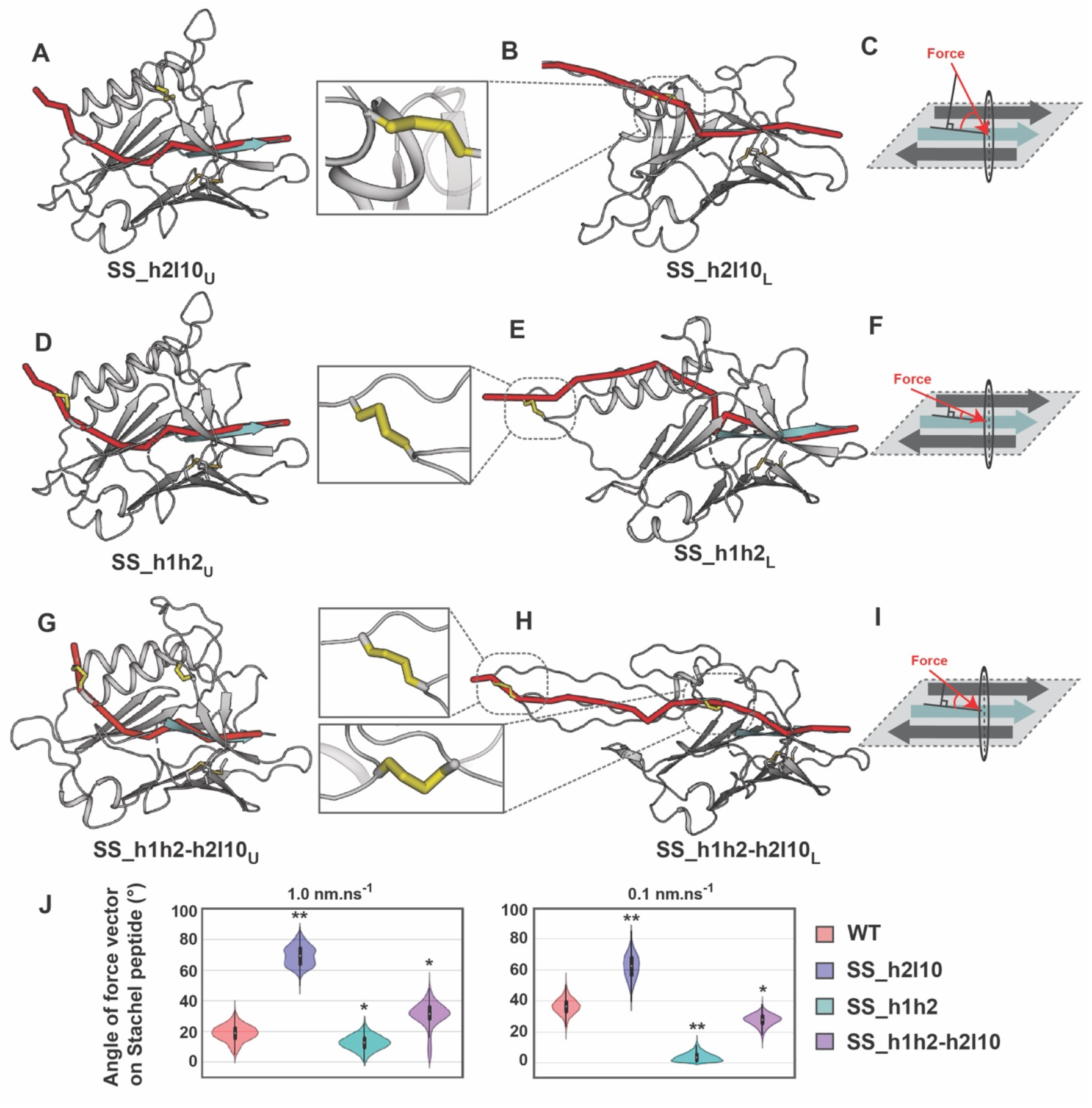
Computational design of GAIN domains with rewired force propagation pathways. (**A-I)** Force propagation path analysis for 3 GAIN variants designed with disulfide bridges locking distinct local regions and rewiring paths (B-D: SS_h2l10; E-G: SS_h1h2; H-J: SS_h1h2-h2l10). From left to right: path representation in the unloaded structure; zoom on designed disulfides; path representation in the loaded structures; schematic representation of the force propagation vector angle reaching the plane constituted by the interacting Stachel and β-strands 6-8. (**J)** Force pathway angle distribution in the fully loaded states of the WT and designed GAIN domains at 0.1 and 1 nm.ns^-1^ pulling speeds (** p-value < 0.001, * p-value < 0.05).

We identified a second variant that prevented rigid body motions within the α-subdomain and locked helices 1 and 2 into the native unloaded conformation through a disulfide bridge (variant referred as SS_h1h2). Consistent with our rationale, the designed GAIN structure did not undergo significant reorientation of the 2 helices in SMD. Although the force propagation path for SS_h1h2 was very similar to WT in the unloaded state (**Fig.3D**), the loaded structure exhibited a different force propagation pathway that traveled through helix 1 to reach the polar motif locking helix 1 onto β-strands 1 and 2, and projected onto the Stachel (**Fig.3E**) with a substantially lower angle (11° and 4° compared to 19° and 36° for the WT optimal path at 1.0 and 0.1 nm.ns^-1^, respectively; **Fig.3F,J**). Due to the increased collinearity of the force vector and rupture interface, we predicted a lower force threshold of GAIN-Stachel dissociation for this variant.

Interestingly, combining the two designed disulfides in the SS_h1h2-h2l10 variant restored WT-like force propagation properties (**Fig.3G-I**). While the pathways run through the two engineered disulfides and travel along loop 10 connecting β-strands 8 and 9, our calculations predicted that the force reaches the Stachel-rupture interface with WT-like vector angle values (31° and 28° at 1.0 and 0.1 nm.ns^-1^, respectively; **Fig.3J**). Overall, despite bearing two designed disulfides that lock and rewire its topology, and should also considerably enhance its thermodynamic stability, the SS_h1h2-h2l10 GAIN was predicted to behave very similarly to WT under mechanical load and release the Stachel at WT-like applied force threshold. Consistent with our path-based predictions, the maximal force resistance observed in our simulations was higher for SS_h2l10 (1450-1600 pN) than for SS_h1h2-h2l10 (1100-1200 pN) and SS_h1h2 (750-950 pN) and suggested distinct mechanical stabilities for the 3 variants.

These 3 computationally designed GAIN variants plus WT represented a unique opportunity to test our understanding of the relationship between topology, structure, dynamics, thermodynamics and mechanical stability that govern the GAIN scaffold. To address these points, we validated experimentally the 3 designed GAINs using structural and biophysical techniques.

To ensure that the engineered disulfide bridges were correctly folded, we attempted to characterize the designed GAIN structures using X-ray crystallography (**Methods**). While the murine ADGRG1 GAIN structure was solved in presence of the ligand adhesion PLL domain and a bound antibody (*27*), we obtained crystals diffracting at 1.9 Å for the human SS_h1h2-h2l10 GAIN construct alone. The structure revealed an overall β-sandwich and α-domain structure similar to that of the murine form (Cα RMSD = 1.2 Å), but diverged significantly in the loop regions owing to significant sequence and structure divergence in these regions (**Fig.4A**). The structure matched our homology models very closely and the conformation of the designed disulfide bridges were superimposable to our predicted bonds (Cα RMSD = 2 Å). Correctly formed disulfide bridges should also enhance the thermodynamic stability of the folded ground state of the protein. Hence, we measured the thermal unfolding of the GAIN domains by circular dichroism and observed enhanced melting temperatures (Tm) for all designs when compared to WT. While the introduction of one disulfide increased the Tm by 4 and 9 °C for SS_h1h2 and SS_h2l10, respectively, the SS_h1h2-h2l10 GAIN reached a Tm as high as 83 °C, (a 17 °C increase from WT) (**Fig.4B**).

**Fig.4.**
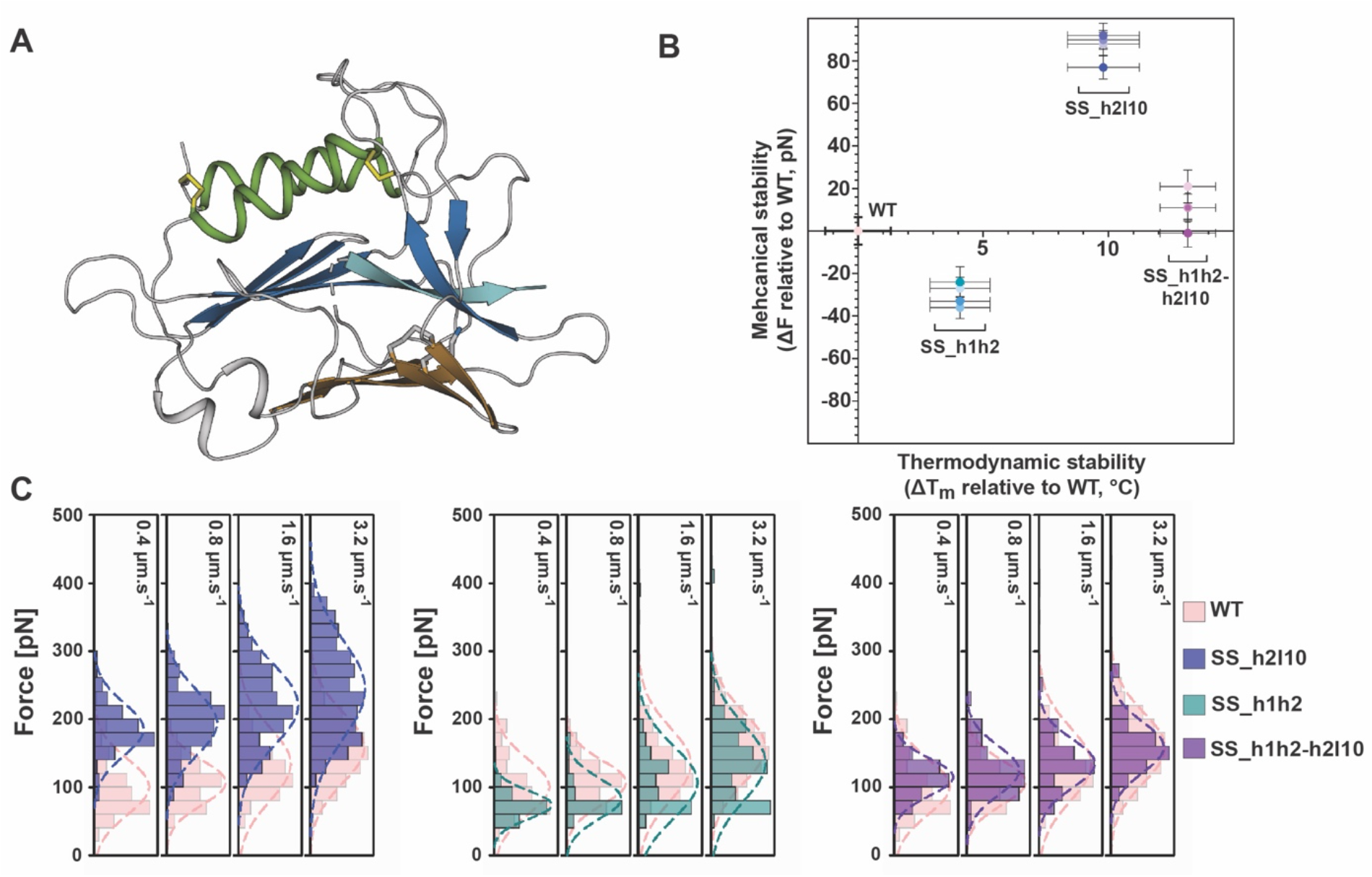
Designed GAIN variants exhibit distinct levels of mechanical stability. (A) X-ray structure of the designed SS_h1h2_h2l10 human ADGRG1 GAIN domain. (**B)** 2D map of the changes in mechanical stability versus thermodynamic stability relative to WT (set at zero, pink dot). **C.** Single-molecule AFM rupture force histograms of the designed GAIN domains at cantilever retraction velocities of 0.4 μm.s^-1^, 0.8 μm.s^-1^, 1.6 μm.s^-1^ and 3.2 μm.s^-1^. A Gaussian fit (dashed line) with the fit parameters are shown for each distribution.

The structural and biophysical characterizations validated the formation of the disulfide bonds, as well as our structural models of the designed protein ground states and prompted us to characterize the mechanical properties of the variants. Following the same functionalization of the protein and experimental conditions as WT, we recorded hundreds of AFM force-extension curves, filtered for single-molecule interactions and quantified GAIN-Stachel dissociation forces. Consistent with our predictions, the Stachel in SS_h1h2 dissociated at significantly lower applied forces than WT (67-135 pN) while the mechanical resistance of the rupture interface increased by almost 2-fold in the SS_h2l10 variant (182-236 pN) (**Fig. 4C**). The combination of disulfides in the SS_h1h2-h2l10 variant restored the typical mechanical resistance observed for the WT domain (115-158 pN, **Fig. 4C**). These data indicated that thermodynamic and mechanical stability do not necessarily correlate with each other as they correspond to distinct properties of the ground and excited states of the protein and can be engineered independently. They also validate our design predictions and suggest that protein mechanics can be engineered by reprogramming long-range dynamic properties of the protein scaffold under mechanical load.

Overall, our results reveal that, GAIN shedding and Stachel dissociation occur primarily under higher mechanical stress than other signaling mechanosensors such as Notch. This key finding suggests a model of ADGRG1 mechanotransduction that involves distinct but complementary activation mechanisms (**Fig. 5**). Low to moderate mechanical stress would primarily induce structural deformations in the ECR disrupting for example non-covalent GAIN-7TM and/or GAIN-PLL interactions, and trigger allosteric signaling responses in the 7TM domain without involving complete Stachel dissociation. Under high mechanical stress, a mechanism of receptor activation through GAIN shedding and Stachel dissociation would predominate and lead to potent signaling responses. The co-existence of both activation modes would enable gradual signaling responses to distinct mechanical perturbations and the integration of other stimuli such as the binding of multiple ligands and co-receptors.

**Fig. 5.**
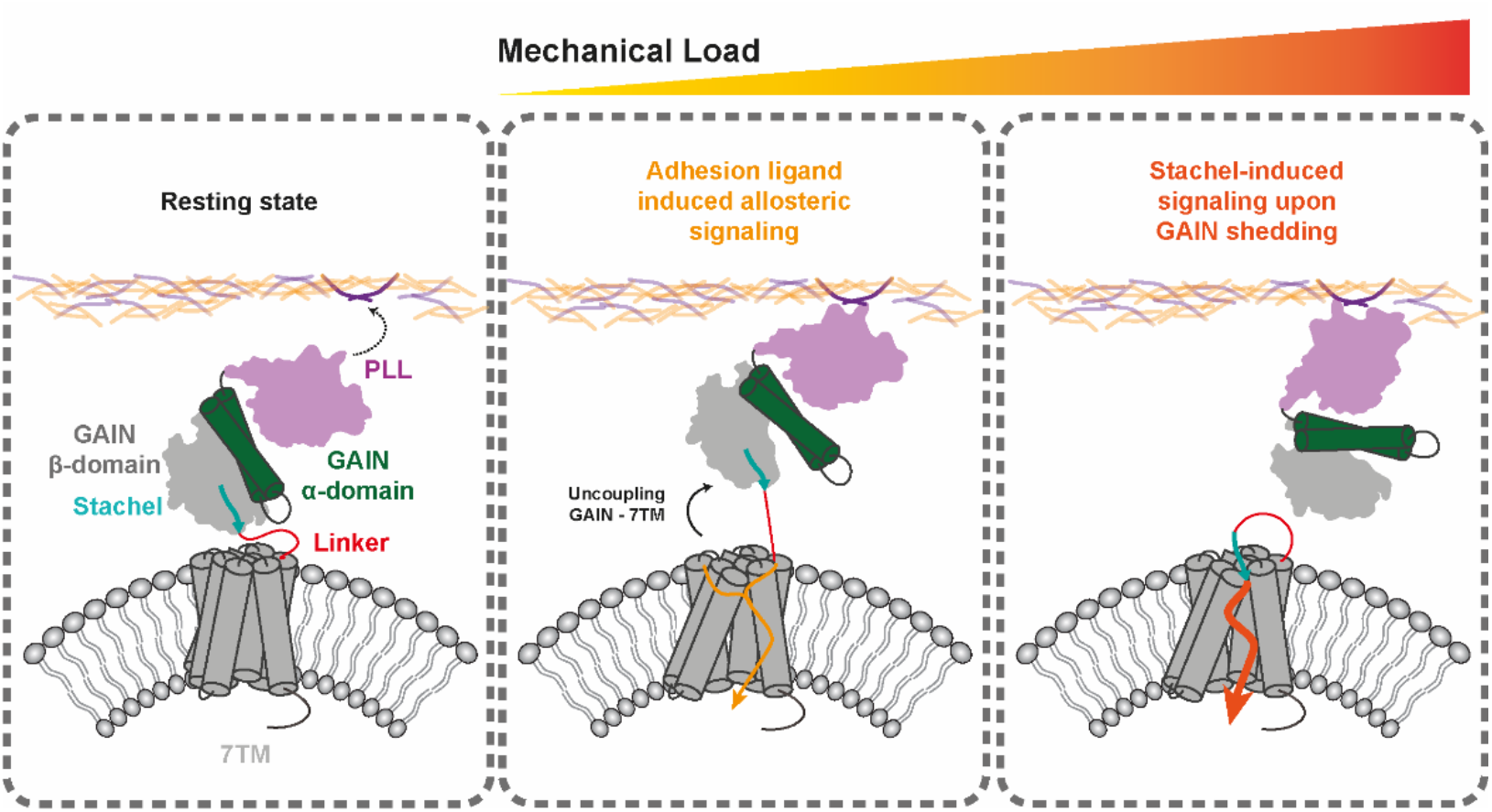
Putative model of ADGRG1 signaling. Schematic description of two distinct but complementary mechanisms of adhesion ligand-induced mechanical activation of ADGRG1. <u>Left.</u> Multi-domain architecture of the full length ADGRG1 binding to ECM ligands through PLL. <u>Middle.</u> Low to moderate levels of mechanical stress trigger structural deformations of the extracellular region including the disruption of non-covalent interactions between GAIN and the PLL and/or 7TM domains. This results in a series of allosteric signals propagating to the 7TM and activating the receptor. <u>Right.</u> High levels of mechanical stress are strong enough to shed the GAIN and release the Stachel which can bind to the 7TM and directly activate the receptor.

While we were preparing this manuscript, two studies were posted to biorxiv (*50*, *51*) that reported measurements of mechanical stability of GAIN domains using single-molecule Magnetic Tweezers experiments. In one of these studies (*51*), the same aGPCR as reported here (ADGRG1) was found to undergo structural changes in a range of 1-20 pN are sufficient to dissociate the Stachel from GAIN and involve significant conformational changes prior to dissociation. While the observation of mechanical induced structural changes is consistent with our simulations, our AFM measurements report higher rupture forces. These discrepancies could be due to significantly different loading rates. Our work analyzes the ADGRG1 over a range from 1,000 – 40,000 pN/sec and when we extrapolate our data to the equivalent loading rate of the other study (1 pN/sec), we obtain values in a range of 20 pN. Another contributing factor could be the stiffness of the potential provided by the magnetic beads is much lower than that of our AFM cantilever. Our data furthermore do not rule out structural transitions in GAIN occurring at force values below that of our fingerprint domains.

## Acknowledgments

We thank Luciano Abriata for help in setting up the CD experiments, the Barth lab members and Alex Persat for insightful discussions and critical comments on the manuscript.

## Author contributions

PB and LD conceived the project. PB supervised the entire project. MN supervised the AFM measurements. FP supervised the protein production and the X-ray crystallography. LD made constructs, purified protein, performed and analyzed CD data. LD and MM carried out the protein modeling and design. MM developed the force propagation pathway analysis and design approaches. MM and MH performed the SMD calculations. BSY carried out and analyzed the AFM measurements. KL, LD and ANL produced the proteins in insect cells. KL, LD and ANL carried out the protein crystallization. FP and KL oversaw the X-ray data collection and structure determination. All authors provided comments on the manuscript.

## Supplementary Information Materials and Methods

### Design of GAIN domain constructs for SMFS-AFM experiments

We designed the GAIN domain constructs for AFM experiments based on the following rationale: It is known that a protein’s mechanical response is highly dependent on the loading point and direction of the applied force (*25*, *28*–*32*). In the context of the full-length ADGRG1 receptor expressed at the cell surface, mechanical force is likely transduced to GAIN by the N-terminal adhesion PLL domain bound to an ECM ligand. To best mimic this native orientation and direction of the applied force in our SMFS experiments, we designed a setup where the C-terminal end of GAIN is attached to a solid surface through an ddFLN4 (i.e., FLN) fingerprint domain while the pulling force is applied to the N-terminal end of the domain (Fig. 1A, Fig. S1). To enable precise repeated measurements of Stachel dissociation by AFM and subsequent quantitative analysis, we modified the cantilever with an SdrG-ddFLN4 fusion. This protein can reversibly bind to the Fgβ peptide that was fused to the N-terminus of the GAIN constructs (Fig. S1). The SdrG-Fgβ complex can withstand very high mechanical forces (over 2 nN) (*23*). After Stachel dissociation from GAIN, the cantilever is clogged with the N-terminal fragment of the GAIN. However, SdrG has a very high off-rate in the absence of force, and dissociates spontaneously from the Fgβ peptide, freeing up the SdrG molecule on the cantilever for binding to new GAIN molecules on the surface. This experimental design enables precise repeated measurements and quantification of Stachel dissociation by AFM by probing fresh GAIN molecules on the surface with each approach-retract AFM cycle.

## Protein Expression and Purification

*AFM handles* – SpyCatcher and SdrG proteins for SMFS were prepared as previously reported (*25*, *52*). Constructed recombinant plasmids pET28a-ybbR-His-ELP(MV7E2)3-FLN-SpyCatcher (Addgene #157674) and pET28a-SdrG-FLN-ELP-His-ybbr (Addgene #168047) were transformed into E. coli BL21 (DE3) strain. Cells were cultured in 5 ml of Luria-Bertani (LB) medium with 50 μg ml^-1^ kanamycin at 37 °C overnight. The culture was transferred to 100 mL of Terrific Broth (TB) medium with 50 μg ml^-1^ kanamycin and cultivated at 37 °C and 200 rpm until an optical density at 600 nm (OD600) of ∼0.8-1.0 was reached. Recombinant protein expression was induced upon addition of 0.5 mM isopropyl-β-D-thio-galactopyranoside (IPTG) and the culture was further incubated at 20 °C and 200 rpm for 18 h. The cells were harvested by centrifugation at 4,000 g for 20 min at 4 °C. The cell pellets were stored at -80 °C until further purification. For purification, the harvested cell pellet was resuspended in a lysis buffer (50 mM Tris, 50 mM NaCl, 0.1% Triton X-100, 5 mM MgCl2; pH 8.0), and disrupted with a sonic dismembrator. The lysate was centrifuged at 14,000 g for 20 min at 4 °C. The supernatant was collected and incubated with Ni-NTA resin (Thermo Fisher Scientific), loaded onto a column, washed with wash buffer (1x PBS buffer with 25 mM imidazole; pH 7.2), and eluted in elution buffer (1x PBS buffer with 500 mM imidazole; pH 7.2). The eluted protein solution was buffer-exchanged and further purified by Superose SEC column, and finally stored in 33% glycerol at -20 °C.

*GAIN domain* – The ADGRG1 GAIN domain was expressed and purified following published protocols (*27*). Insect codon-optimized human ADGRG1 ECR sequences (Uniprot: Q9Y653) were synthesized (Twist Biosciences) as part of larger constructs for either AFM or X-ray crystallography experiments (see full protein sequences) in a pFastBac vector backbone with a BiP secretion signal. A baculovirus expression system was used for expression of proteins in High-Five insect cells. pFastBac constructs were transformed into DH10MultiBac cells (Geneva Biotech), white colonies indicating successful bacmid recombination were selected, and bacmids were purified by the alkaline lysis method. Sf9 cells were transfected using the X-tremeGENE 9 transfection reagent (Sigma) with the desired bacmid. eYFP-positive cells were observed after 1 week and subjected to one round of viral amplification. Amplified P2 virus was used to infect High-Five cells at a density between 1–2 × 10^6^ cells/ml. Cells were incubated for 72 h at 28°C, pelleted (1000 x *g*, 10 minutes) and the recovered supernatant was clarified (5000 x *g*, 30 minutes) and filtered using a Grade GD 1 μm pre-filter (1841-047, Whatman, GE) on top of a Durapore 0.45 μm PVDF Membrane filter (HVLP04700, Merck Milipore). 5 ml of 50% Fastback Ni Advance Resin slurry (Protein Ark) pre-equilibrated in HBS buffer containing 10 mM HEPES (pH 7.2) and 150 mM NaCl was added to the 500 ml clarified supernatant containing the secreted proteins and incubated over-night at 4°C on a rotating wheel. The resin was recovered on a gravity-flow column, equilibrated with 30 ml HBS and washed with 30 ml HBS containing 30 mM imidazole. Proteins were eluted with 15 ml of HBS containing 300 mM imidazole and 1 ml fractions were collected and analyzed by SDS-PAGE. The three or four purest and most concentrated fractions were pooled and concentrated on a 30 kDa Amicon Ultra-4 centrifugal filter (UFC803024, Merck Millipore) down to 500 μl. The concentrated proteins were loaded on a Superdex 200 Increase 10/300 GL (28-9909-44, Sigma-Aldrich) connected to an ÄKTA Pure chromatography system (GE) and eluted in HBS. The fractions containing the peak corresponding to the purified GAIN construct were pooled and quantified using A_280_ absorbance. Purified GAIN proteins were either directly concentrated to 10 mg.ml^-1^ for crystallization or flash-frozen in liquid nitrogen and stored at -80°C for subsequent AFM experiments.

### AFM-SMFS surface preparation and protein immobilization

The surface modification of cantilever and coverglasses and the protein immobilization were done as described previously (*52*). Cantilevers were cleaned by UV-ozone treatment for 40 min and coverglasses were soaked in piranha etching solution and rinsed with distilled water (DW). Then, cantilevers and coverglasses were treated with 3-Aminopropyl (diethoxy) methylsilane (APDMES, ABCR GmbH, Karlsruhe, Germany) to silanize the surfaces with amine groups. The amine groups subsequently reacted to a NHS group from sulfosuccinimidyl 4-(N-maleimidomethyl)cyclohexane-1-carboxylate (sulfo-SMCC; Thermo Fischer Scientific) in 50 mM HEPES buffer pH 7.5 for 30 min. The thiol group from Coenzyme A (CoA, 200 μM) reacted to a maleimide group from sulfo-SMCC in coupling buffer (50 mM sodium phosphate, 50 mM NaCl, 10 mM EDTA, pH 7.2) for 2 hrs. Finally, the ybbR-tagged proteins SdrG-FLN-ELP-His-ybbr and pre-conjugated ybbR-His-ELP(MV7E2)3-FLN-SpyCatcher-SpyTag-GAIN variants were site-specifically anchored to the surface using SFP-mediated ligation to CoA in Mg2+ supplemented 1x PBS buffer. This resulted in covalent immobilization of SdrG and GAIN variants to cantilever and cover glasses, respectively. Protein-immobilized cantilevers and coverglasses were extensively washed and kept in 1x PBS buffer prior to immediate use. SpyTag-SpyCatcher conjugation of ybbR-His-ELP-FLN-SpyCatcher and Fgβ-GAIN-His-SpyTag variants were done by mixing two proteins with same molar ratio in PBS buffer and pre-incubation for 1-12 hrs prior to ybbR tag ligation.

### AFM-SMFS measurements and data analysis

Force spectroscopy measurements with SdrG and Fgβ-fused GAIN variants were conducted in the same manner as previously illustrated using automated AFM-based SMFS (Force Robot 300, JPK Instruments). SMFS data were recorded in 1x PBS at room temperature with constant pulling speeds of 400, 800, 1600 and 3200 nm.s^-1^. A total of 258,044 force-extension curves were acquired. Force-extension curves were filtered and analyzed by a combination of software available on the AFM instrument and custom python scripts. The majority of data traces contained no interactions, non-specific interaction, or complex multiplicity of interactions. Therefore, the data traces were filtered by searching for contour length increments that matched the lengths of the fingerprint domains, FLN (≈36 nm). Theoretical contour length increment was calculated based on the equation ΔL_c_ = (0.365 nm/AA) × (# AAs in POI) - L_f_, where ΔL_c_ is expected contour length increment and L_f_ is end-to-end length of folded protein domain. For FLN, ΔL_c_ = 36.9 nm - L_f_, where L_f_ is typically < 5nm. The total number of force-extension curves matching this criterion was 509 out of 120,780, 1,058 out of 68,376, 263 out of 42,295, and 158 out of 18,447 for GAIN-WT, SS_h2l10, SS_h1h2, and SS_h1h2-h2l10, respectively.

For dynamic force spectra, the rupture or unfolding forces vs. loading rate was plotted and median forces and loading rates for each pulling speed were fitted to Bell-Evans model (*53*, *54*) to estimate the effective distance to the transition state (Δx) and the intrinsic dissociation rate or unfolding rate (k_off_) in the absence of force. Data were fitted using the Dudko-Hummer-Szabo model (*55*, *56*) with a cusp-like barrier (𝜈 = 0.5) to estimate Δx, k_off_, and energy barrier height (ΔG).

For direct comparison with the same cantilever, both GAIN-WT and GAIN(4Cys) were immobilized on different areas of the same surface and a single cantilever was used to probe each spot using constant speed pulling at 3200 nm s-1. After 500 approach-retraction cycles at one area, the surface was moved to the other area. This cycle was done once for each GAIN-WT and GAIN(4Cys). The total number of force-extension curves matching the criterion was 69 out of 9,484, and 67 out of 4,986 for GAIN-WT and GAIN(4Cys), respectively.

For contour length analysis in reducing condition, each GAIN-WT and noGAIN was probed with the same cantilever both in non-reducing condition (0 mM DTT in 1x PBS) and reducing condition (100 mM DTT in 1x PBS). SMFS data were recorded in 1x PBS first at room temperature with constant pulling speed of 800 nm s-1, and then buffer was changed to 100 mM DTT in 1x PBS and SMFS data were recorded with the same pulling speed. The total number of force-extension curves matching the criterion was 101 out of 1,566, and 116 out of 26,097 for GAIN-WT and GAIN(4Cys), respectively.

### Circular Dichroism measurements

Purified GAIN variant proteins were adjusted to 10-15 μM final concentration in 10 mM HEPES, 15 mM NaCl, pH 7.5. Thermal denaturation CD spectra were recorded on a Chirascan CD spectrometer (AppliedPhotophysics) from 20°C to 96°C using a 2°C step, between 215 nm and 250 nm. Each melting curve was normalized between 0 and 1 across the entire spectral range and the 216 nm and 232 nm normalized spectra from 2 or 3 replicate experiments were extracted and plotted.

### X-ray crystallization and data collection

Crystals of the GAIN_SS_h2l10-h1h2 variant were grown in sitting drops using the random microseed screening (*57*). (1) 10 mg/mL of purified protein was mixed 1:1 (200 nL) with mother liquor and crystals appeared after several days at 18°C in a buffer containing 0.1 M potassium thiocyanate and 30% PEG 2000 MME. Crystals were harvested and cryoprotected in mother liquor supplemented with 25% glycerol and frozen under liquid nitrogen.

All datasets were collected at beam line PX-III of the Swiss Light Source, Villigen, Switzerland. Datasets were collected with λ = 1.00 Å. All datasets were processed with XDS. (2) The structure was solved by molecular replacement the program Phaser (3), using PDB 5KVM as the initial search model. The final structures were determined after iterative rounds of model-building in COOT (4) and refinement in phenix.refine (5). The model was refined to a R/Rfree of 0.196/0.241 with excellent geometry (no Ramachandran or Rotamer outliers). Model was validated with Molprobity, with a score of 1.26. (6) Final statistics were generated as implemented in phenix.table_one (*57*–*62*).

### Computational variant design

To screen for possible pairs of residues enabling the formation of disulfides, the Coupled Moves algorithm from the Rosetta software was used with the backrub movers protocol (*63*). A total of 100 independent structures was generated for every pair and the top 10% lowest energy structures were selected. All structures were then minimized and relaxed over all conformational degrees of freedom to reach the lowest energy minimum conformation through the relax protocols included in the Rosetta package. Distances between the sulfide groups of added cysteines were computed and averaged across the lowest energy structures. To determine which pairs of mutations could form a disulfide, a distance cutoff of 2.15 Å between the 2 cysteines was used to conserve or exclude mutants.

### SMD simulations

The structure of the GAIN_SS-h2l10-h1h2 variant was resolved at 1.92 Å resolution and back-mutated using the above protocol to recover the individual disulfide variants as well as the WT structure. The system was solvated with a box size of 80 Å along X, Y and Z dimensions with 0.1 M of 𝑁𝑎^+^ and 𝐶𝑙^-^ ions. The total system size was approximatively 130.000 atoms. Simulations were performed with GROMACS 2019.4 (*64*) using the CHARMM36 forcefield (*65*) along with the TIP3 water model (*66*) in an *NPT* ensemble at 310K and 1 bar using a velocity rescaling thermostat (with a relaxation time of 0.1 𝑝𝑠) and Berendsen barostat (with a relaxation time of 1 𝑝𝑠). Equations of motion were integrated with a timestep of 2 𝑓𝑠 using a leap-frog algorithm. Each system was energy minimized using a steepest descent algorithm for 5000 steps, and then equilibrated with the atoms of the protein restrained using a harmonic restraining force in two steps, first by adding the thermostat for 50000 steps and then the barostat for 500000 steps. SMD simulations were performed using the umbrella sampling method with a pulling speed ranging from 1 Å/ns to 10 Å/ns. In all simulations, the 2 alpha carbons of the N-term and C-term residues were moved harmonically in the desired direction with constant velocity. A distance cutoff of 14 Å was used for short-range non-bonded interactions, whereas electrostatic interactions were computed using the Particle-mesh Ewald method (*67*) with a cubic interpolation of power 4.

---

**Fig. S1.**
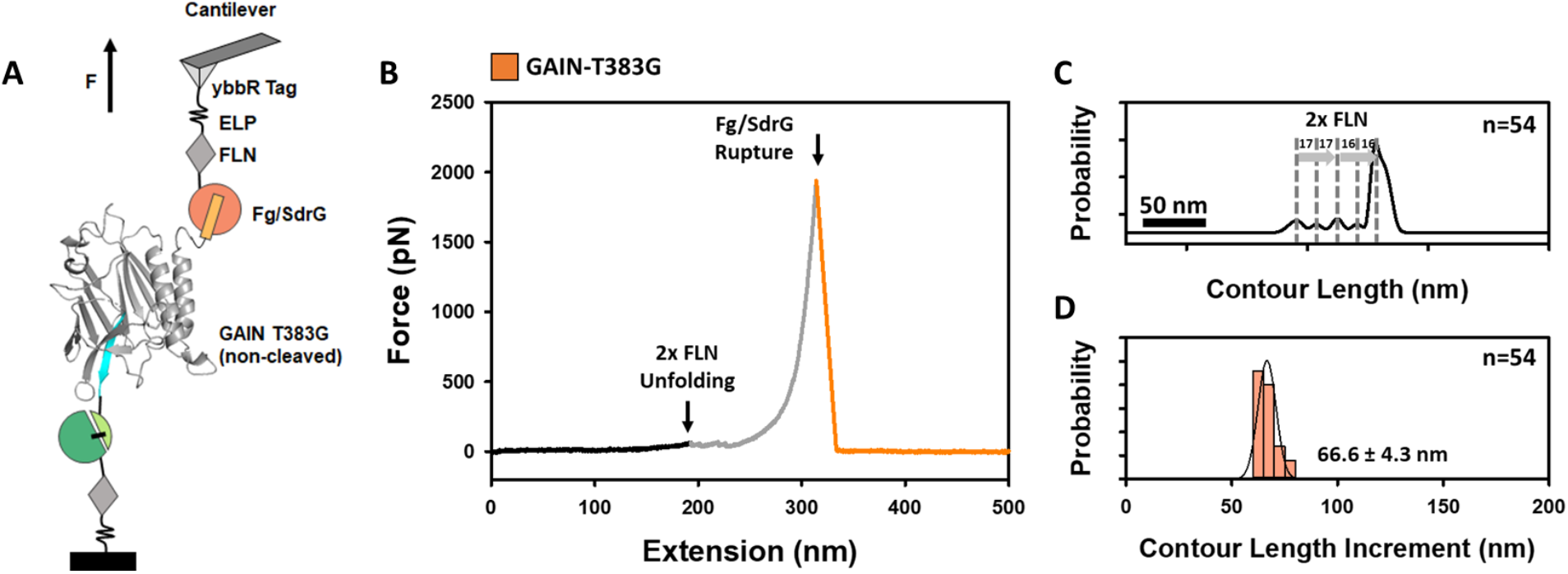
AFM-SMFS of the uncleavable GAIN. (A) Schematic illustration of experimental configuration with uncleavable GAIN-T383G. (B) Typical force-extension trace. Unfolding of two FLN fingerprint domains (gray) was used to filter the curves for specific single-molecule interactions. Fingerprint unfolding was followed by Fg:SdrG complex rupture (orange). (C) Contour length analysis (n = 54). (D) Contour length increment histogram (n = 54). Two ∼34 nm contour length increments with intermediate folding states associated with FLN unfolding were observed. Additional force peak or contour length increase was not observed. Measurement and analyses were done with a pulling speed of 800 nm s^−1^.

**Fig. S2.**
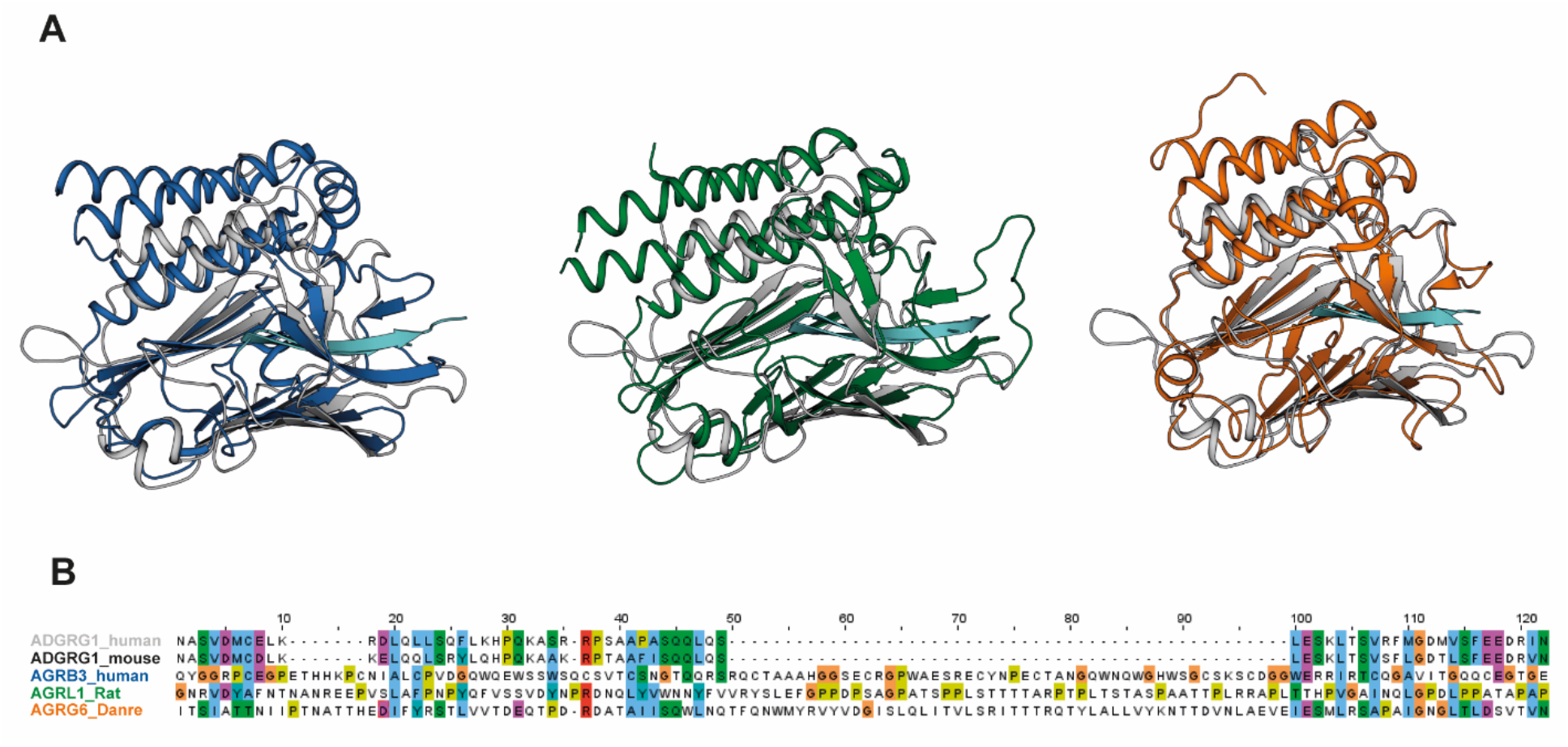
Structural comparison of our modelled human ADGRG1 GAIN domain with other known GAIN structures. (A) Structural alignment of the GPR56 GAIN domain (gray) reconstructed from the crystal structure of the SS_h1h2-h2l10 variant with the crystal structure of other GAIN domains (4DLO: blue, 4DLQ: green, 6V55: orange). (B) Sequence alignment of the alpha helical subdomain of the human GAIN domain with the other GAIN domains.

